# Genome-wide identification of genes important for growth of *Dickeya dadantii* and *D. dianthicola* in potato (*Solanum tuberosum*) tubers

**DOI:** 10.1101/2021.09.16.460530

**Authors:** Tyler C. Helmann, Melanie J. Filiatrault, Paul V. Stodghill

**Affiliations:** Emerging Pests and Pathogens Research Unit, Robert W. Holley Center for Agriculture and Health, Agricultural Research Service, United States Department of Agriculture, Ithaca, New York, USA; School of Integrative Plant Science, Plant Pathology and Plant-Microbe Biology Section, Cornell University, Ithaca, New York, USA

**Keywords:** potato, soft rot, RB-TnSeq, TnSeq, *Dickeya dadantii*, *Dickeya dianthicola*

## Abstract

*Dickeya* species are causal agents of soft rot diseases in many economically important crops, including soft rot disease of potato (*Solanum tuberosum*). Using random barcode transposon-site sequencing (RB-TnSeq), we generated genome-wide mutant fitness profiles of *Dickeya dadantii* 3937, *Dickeya dianthicola* ME23, and *Dickeya dianthicola* 67-19 isolates collected after passage through several *in vitro* and *in vivo* conditions. Tubers from the potato cultivars “Atlantic”, “Dark Red Norland”, and “Upstate Abundance” provided highly similar conditions for bacterial growth. Using the homolog detection software PyParanoid, we matched fitness values for orthologous genes in the three bacterial strains. Direct comparison of fitness among the strains highlighted shared and variable traits important for growth. Bacterial growth in minimal medium required many metabolic traits that were also essential for competitive growth *in planta*, such as amino acid, carbohydrate, and nucleotide biosynthesis. Growth in tubers specifically required the pectin degradation gene *kduD*. Disruption in three putative DNA-binding proteins had strain-specific effects on competitive fitness in tubers. Though the Soft Rot *Pectobacteriaceae* can cause disease with little host specificity, it remains to be seen the extent to which strain-level variation impacts virulence.

## Introduction

The Soft Rot *Pectobacteriaceae* comprise *Dickeya* and *Pectobacterium* species that are the causal agents of bacterial soft rot diseases on economically-important vegetables and ornamentals (Adeolu et al., 2016; Motyka et al., 2017). These necrotrophic pathogens rely on plant cell wall degrading enzymes (PCWDEs) to form visible symptoms, as well as numerous traits to survive conditions encountered in the host such as oxidative stress, osmotic stress, iron starvation, and toxic compounds (Jiang et al., 2016; Reverchon et al., 2016). The taxonomy within these genera has undergone substantial revision with the addition of novel species *D. solani, D. aquatica*, and *D. fangzhongdai* (Samson et al., 2005; Parkinson et al., 2014; Wolf et al., 2014; Tian et al., 2016). However, an increase in available whole-genome sequence data has improved species-level identification based on pairwise average nucleotide identity (ANI), *in silico* DNA-DNA hybridization (*is*DDH), and core genome multilocus sequence analysis (MLSA) (Zhang et al., 2016). There is little known about host-specific traits, as these species generally have broad host ranges (Van Gijsegem et al., 2021). In addition, there are no known resistance genes for potato soft rot, and it is therefore impossible to predict cultivar resistance without testing (Lyon, 1989; Czajkowski et al., 2011; Chung et al., 2013). Without gene-for-gene resistance, potato cultivar tolerance is reliant on physical barriers and antimicrobial small molecules such as phenolics or the phytoalexin rishitin (Lyon, 1989). An alternative strategy being explored is the use of bacteriophage-based biocontrol for potato plants and tubers, particularly of the highly-virulent *D. solani* (Adriaenssens et al., 2012; Czajkowski et al., 2017).

*Dickeya* virulence factors and transcriptional regulators of virulence genes are generally conserved. Studies in *D. solani* have suggested a closed pangenome with many conserved virulence factors and transcriptional regulators (Golanowska et al., 2018; Motyka-Pomagruk et al., 2020). However, virulence regulon differences indicate some virulence genes could have differential expression among strains (Golanowska et al., 2018). Pangenomic analysis of *D. dianthicola* also reflects a closed pangenome, though almost all sequenced strains were originally isolated from potato (Ge et al., 2021b).

To identify bacterial traits important for growth in potato (*Solanum tuberosum*) tubers, we examined three strains across two species: *D. dadantii* 3937 (*Dda*3937), *D. dianthicola* ME23 (*Ddia*ME23), and *D. dianthicola* 67-19 (*Ddia*6719). While these three strains are all pathogenic on potato, *Dda*3937 was originally isolated from *Saintpaulia ionantha* (Lemattre and Narcy, 1972), and *Ddia*6719 was originally isolated from New Guinea impatiens (*Impatiens hawkeri*) (Liu et al., 2020a, 2020b). *Dda*3937 has been a model strain used for molecular studies since its isolation in 1972 (Lemattre and Narcy, 1972), while *Ddia*ME23 was isolated as a representative strain for a 2014 potato disease outbreak (Ma et al., 2019). Pairwise ANI between *Dda*3937 and *Ddia*ME23 is 92.8% (Chen et al., 2019). Type strain *D. solani* IPO 2222 has a pairwise ANI score of 94.7% to *D. dadantii* 3937 (Chen et al., 2019). We aimed to directly test whether there was any variation in the contributions to competitive fitness for virulence traits and regulators.

Transposon mutagenesis of *P. carotovorum* followed by screening for altered soft rot symptoms in Chinese cabbage identified genes involved in nutrient utilization, production of PCWDEs, motility, biofilm formation, and toxin susceptibility (Lee et al., 2013). Transposon mutagenesis followed by high-throughput sequencing (TnSeq) is a valuable screening tool to identify genes important for growth in a given condition (van Opijnen et al., 2009). TnSeq has been used to identify *D. dadantii* genes important for growth in chicory (Royet et al., 2019). This work identified several metabolic pathways essential for *in planta* growth, primarily those involved in biosynthesis of nucleotides, amino acids, and some vitamins (Royet et al., 2019). A modification of TnSeq to add 20-nucleotide DNA “barcodes” to transposon donor plasmids, known as random barcode transposon-site sequencing (RB-TnSeq) enables highly-scalable TnSeq assays (Wetmore et al., 2015). This method has been applied to over 44 bacterial strains to date (Price et al., 2018), including plant pathogenic *Pseudomonas* spp. and *Ralstonia* spp. (Cole et al., 2017; Helmann et al., 2019; Georgoulis et al., 2020). By leveraging RB-TnSeq in a shared susceptible host for *D. dadantii* and *D. dianthicola*, we aimed to identify common and unique virulence factors among representative strains for these two species.

## Materials and Methods

### PyParanoid gene ortholog group assignments

Gene ortholog groups were generated using the PyParanoid analysis pipeline v0.4.1 (Melnyk et al., 2019). Peptide sequences from the following RefSeq genome assemblies were used to construct ortholog groups: GCF_000147055.1 (*Dda*3937), GCF_003403135.1 (*Ddia*ME23), and GCF_014893095.1 (*Ddia*6719). From these assemblies, RefSeq gene loci were then matched to their corresponding protein names to allow comparison to the Barcode Sequencing (BarSeq) fitness data. Additionally, Clusters of Orthologous Groups (COG) categories for *Dda*3937 were downloaded from the IMG database (Chen et al., 2019), GenBank gene names were replaced with their corresponding RefSeq names, and added to this ortholog table, allowing putative COG assignments for orthologous genes in *D. dianthicola* strains.

### Barcoded transposon library construction

Strains used in this study are described in Table S1. All bacteria were cultured in Luria-Bertani (LB) medium (10g tryptone, 5g yeast extract, and 10g NaCl per 1L) (Bertani, 1951) at 28°C. Barcoded transposon libraries were constructed as previously described (Wetmore et al., 2015). Briefly, barcoded *mariner* transposon plasmid pKMW3 was conjugated from the *E. coli* WM3064 donor library APA752 (Wetmore et al., 2015) into wild-type *Dda*3937, *Dia*ME23, and *Ddia*6719, each on 50 LB plates containing 300µM diaminopimelic acid (Sigma-Aldrich, USA). Conjugations were incubated at 28°C overnight, and exconjugants were then scraped into 10mM KPO_4_. This conjugation mixture was then spread onto 200 LB plates per strain, containing 50µg/ml kanamycin and incubated at 28°C for 3 days. All colonies were resuspended in 200ml LB with kanamycin, diluted to OD_600_ 0.2, and grown at 28°C with shaking until OD_600_ reached 1.5 – 3.0, which corresponded to approximately 6 to 8 hours. Glycerol was added to the library to a final concentration of 15%, and 1ml aliquots were frozen at -80°C.

### DNA library preparation and sequencing

For DNA library preparation, genomic DNA from each library was purified from an entire 1ml cell pellet using the Monarch Genomic DNA Purification Kit (New England Biolabs, USA). Samples were eluted in 50µl nuclease-free water. Purified DNA was quantified on a Nanodrop One (Thermo Fischer Scientific, USA), and 500ng DNA was used as input for the NEBNext Ultra II FS DNA Library Prep kit (New England Biolabs, USA), following the manufacturer’s instructions with modifications as follows. For enzymatic DNA fragmentation, a 12-minute incubation time was used. DNA fragments were size selected using AMPure XP magnetic beads (Beckman Coulter, USA) at the recommended ratios 0.4X and 0.2X. We used a modified version of the protocol described in (Wetmore et al., 2015), with a two-step PCR used to enrich for transposon insertion sites, based on (Rubin et al., 2020). A custom splinkerette adapter was ligated to fragmented DNA, prepared by annealing oligos: /5Phos/G*ATCGGAAGAGCACACGTCTGGGTTTTTTTTTTCAAAAAAA*A and G*AGATCGGTCTCGGCATTCCCAGACGTGTGCTCTTCCGATC*T (Rubin et al., 2020). Between rounds of PCR and before submitting for sequencing, DNA was cleaned by binding to AMPure XP magnetic beads, using a bead ratio of 0.9X and eluted in 15µl 0.1X TE buffer for intermediate steps and 30µl 0.1X TE for sequencing. Finally, the sequencing library was quantified using a Qubit dsDNA HS assay kit (Thermo Fischer Scientific, USA). DNA libraries were submitted for sequencing at the Biotechnology Resource Center (BRC) Genomics Facility at the Cornell Institute of Biotechnology on an Illumina MiSeq to check library quality, followed by sequencing on a NextSeq 500 (Illumina, Inc. USA). All mapping used single-end sequencing for 150bp fragments.

### Transposon library mapping

Sequence data were analyzed using the scripts MapTnSeq.pl and DesignRandomPool.pl from the FEBA package v1.3.1 (Wetmore et al., 2015) to map reads to the genome and assemble the mutant pool using barcodes seen in a single location 10 or more times. The transposon sequence “model_pKMW3.2” was used to identify transposon sequence in the reads. All TnSeq mapping and BarSeq fitness calculation code is available at http://bitbucket.org/berkeleylab/feba/ (Wetmore et al., 2015). Mapping scripts were run on a Cornell University BioHPC Cloud 40-core Linux (CentOS 7.6). server with 256GB RAM.

### Gene essentiality predictions

Using the output from MapTnSeq.pl, gene essentiality predictions were made using https://bitbucket.org/berkeleylab/feba/src/master/bin/Essentiality.pl and the function “Essentials” from https://bitbucket.org/berkeleylab/feba/src/master/lib/comb.R (Wetmore et al., 2015). Using the median insertion density and the median length of genes >100bp, this method calculates how short a gene can be and still be unlikely to have no insertions by chance (P <0.02, Poisson distribution); genes shorter than this threshold are then excluded (Price et al., 2018). For the *Dickeya* strains examined here, the minimum gene length for a gene to be predicted as essential for growth in LB was 175bp (*Dda*3937) or 150bp (*Ddia*ME23 and *Ddia*6719). Protein-coding genes are then considered to be essential or nearly-essential if there are no fitness values and the normalized central insertion density score and normalized read density score as computed by the FEBA package were <0.2 (Price et al., 2018).

### Library pre-culture

For a given BarSeq experiment, a single transposon library freezer aliquot was thawed and recovered in 25ml LB containing 50µg/ml kanamycin at 28°C until OD600 ∼ 0.5 to 0.7, approximately 6 to 8 hours. At this point, two 1ml cell pellets were frozen as time0 measurements, and the remaining culture was washed in 10mM KPO_4_ and used to inoculate experimental samples.

### *In vitro* samples

All *in vitro* cultures were grown in 1ml volumes in 24-well plates. In each well, 50µl starter culture at 0.3 OD600 was added to 950µl medium containing 50µg/ml kanamycin. Media tested were LB, Potato Dextrose Broth (PDB) (Sigma-Aldrich, USA), and M9 minimal medium (M9) as described in (M9 minimal medium (standard), 2010) but containing 0.4% glycerol instead of 0.4% glucose. Plates were incubated at 28°C with shaking at 200 rpm. After 1 day (LB and PDB) or 2 days (M9), each 1ml sample was pelleted and frozen prior to genomic DNA extraction.

### Tuber samples

Prior to inoculation, all tubers were rinsed and then surface sterilized by submerging in 70% ethanol for 10 minutes, followed by two washes with distilled water. Inoculum was standardized to OD600 3.0 (approximately 10^9^ CFU/ml), and 10µl was inoculated in two replicate stab wounds created by pushing a 200µl pipet tip roughly 3mm into each tuber. Six replicate tubers were used for each bacterial strain and potato cultivar. Inoculated tubers were stored in plastic bags at 28°C. Two days after inoculation, ∼2cm length cores were taken at each site of inoculation using a 1cm diameter cork borer. Duplicate cores from each tuber were pooled in 8ml 10mM KPO_4_ and shaken at 200rpm at 28°C for 10 minutes. For each sample, 2ml bacterial suspension was pelleted and frozen prior to DNA extraction.

### BarSeq PCR and sequencing

Genomic DNA was purified from cell pellets using the Monarch Genomic DNA Purification Kit (New England Biolabs, USA). Purified DNA samples were eluted in 30µl nuclease-free water and quantified on a Nanodrop One (Thermo Fischer Scientific, USA). After gDNA extraction, the 98°C BarSeq PCR as described in (Wetmore et al., 2015) was used to specifically amplify the barcode region of each sample. The PCR for each sample was performed in 50µl total volume: containing 0.5µl Q5 High-Fidelity DNA polymerase (New England Biolabs, USA), 10µl 5X Q5 buffer, 10µl 5X GC enhancer, 1µl 10mM dNTPs, 150 to 200 ng template gDNA, 2.5µl common reverse primer (BarSeq_P1), and 2.5µl of forward primer from one of the 96 indexed forward primers (BarSeq_P2_ITXXX), both at 10µM (Wetmore et al., 2015). Following the BarSeq PCR, 10µl of each reaction was pooled (46 to 49 samples per pool), and 200µl of this DNA pool was subsampled and purified using the DNA Clean & Concentrator Kit (Zymo Research, USA). The final DNA sequencing library was eluted in 30µl nuclease-free water, quantified on a Nanodrop One, and submitted for sequencing at the BRC Genomics Facility at the Cornell Institute of Biotechnology. Each sequencing pool was run on a single NextSeq 500 (Illumina, Inc., USA) lane for 75 bp single-end reads.

### Gene fitness calculations

Sequencing reads were used to calculate genome-wide gene fitness using the FEBA scripts MultiCodes.pl, combineBarSeq.pl, and BarSeqR.pl (Wetmore et al., 2015). Scripts to calculate gene fitness values were run on a Cornell University BioHPC Cloud 40-core Linux (CentOS 7.6). server with 256GB RAM. Fitness values for each gene were calculated as the log_2_ ratio of relative barcode abundance following library growth under a given condition divided by the relative abundance in the time0 sample. Barcode counts were summed between replicate time0 samples. For analysis, genes were required to have at least 3 reads per strain and 30 reads per gene in the time0 sample (Wetmore et al., 2015). The fitness values were calculated based on the “central” transposon insertions only, i.e., those within the central 10% to 90% of a gene. The normalized median gene fitness value was 0. All experiments described here passed previously described quality control metrics (Wetmore et al., 2015).

### Fitness analysis and plotting

We focused on genes having fitness values >1 or <-1 and absolute t-like test statistic >4. This t score is an estimate of the reliability of the fitness measurement for a gene, and is equal to the fitness value divided by the square root of the maximum variance calculated in two ways (Wetmore et al., 2015). With these cutoffs, we also calculated gene fitness values comparing replicate time0 samples (Price et al., 2018; Liu et al., 2021). Across 6 (each *Dda*3937 and *Ddia*ME23) and 2 (*Ddia*6719) replicate time0 samples, 0 gene fitness values had fitness >1 or <-1 and absolute t >4. Data were analyzed in R v4.0.3 (R Core Team, 2017) using the package ggplot2 v3.3.5 (Wickham, 2016). The principal components analysis was performed on the gene fitness matrix for each strain using the R function prcomp, which performs centered singular value decomposition.

### Data availability Statement

All raw Illumina reads used for mapping and fitness assays have been deposited in the Sequence Read Archive under BioProject accession number PRJNA692477. Individual sample accession numbers are listed in Table S2. Annotated scripts used for computational analysis are available at github.com/tylerhelmann/dickeya-barseq-2021. Experimental fitness values are publicly available at fit.genomics.lbl.gov.

## Results

### Identification of homologous gene families in *D. dadantii* 3937, *D. dianthicola* ME23 and *D. dianthicola* 67-19

To enable direct comparison of gene fitness measurements between strains, we constructed a database of homologous gene families using the PyParanoid analysis pipeline (Melnyk et al., 2019). Based on clustering of all predicted protein sequences from *Dda*3937, *Ddia*ME23, and *Ddia*6719, 3,821 total homolog groups were identified, representing 88.1% of the total input sequences. Of these, 3,310 groups contained single-copy genes in all three strains. For each group, gene loci, protein identifiers, and gene descriptions are listed in Table S3. This table also contains COG assignments matched from the *Dda*3937 IMG genomic annotation (Chen et al., 2019).

### Creation of barcoded transposon libraries in *Dickeya* spp

To measure contributions of individual genes to fitness, we constructed barcoded transposon mutant libraries in the *Dickeya* strains using a barcoded *mariner E. coli* donor library (Wetmore et al., 2015). These libraries ranged in size from 334,893 to 541,278 mapped genomic insertional strains, with 37 to 62 median strains per hit gene (Table 1). Of the three strains tested, only one gene in the *Ddia*6719 genome did not contain any TA dinucleotide sites and was therefore inaccessible to the *mariner* transposon. Mapped insertions were evenly distributed across the chromosome of each strain (Fig. S1).

**Table 1.** Characteristics of the *mariner* transposon libraries. “Central” insertions are those within the central 10 – 90% of a gene.

| Library | Insertions in genome | Central insertion strains | Genes with central insertions (Total) | Median strains per hit gene |
| --- | --- | --- | --- | --- |
| *D. dadantii* 3937 | 337,541 | 193,696 | 3,882 (4,213) | 37 |
| *D. dianthicola* ME23 | 541,278 | 321,087 | 3,805 (4,182) | 62 |
| *D. dianthicola* 67-19 | 334,893 | 200,170 | 3,728 (4,110) | 41 |

### Identification of essential gene sets in *D. dadantii* and *D. dianthicola*

Based on analysis of the TnSeq mapping data, essential genes were predicted using the FEBA RB-TnSeq analysis pipeline (Wetmore et al., 2015; Price et al., 2018). We identified 374 to 426 genes per strain that are likely to encode essential or near-essential genes for growth in LB (Table S4). Using the ortholog group assignments for these genes, 316 of these predicted essential genes (74 to 84%) are shared among all three strains (Fig. S2). Most predicted essential genes are in the functional categories of “Translation, ribosomal structure, and biogenesis”, “Cell wall/membrane/envelope biogenesis”, “Coenzyme transport and metabolism”, “Energy production and conversions”, and “Replication, recombination, and repair” (Table S5).

### Conducting pooled growth assays to measure relative mutant fitness

To generate genome-wide gene fitness values for the barcoded transposon libraries, each strain was grown in the rich media LB and Potato Dextrose Broth (PDB) as well as M9 minimal medium supplemented with 0.4% glycerol (Fig. S3). Strain fitness values were calculated as a log2 ratio of barcode abundance following sample growth with barcode abundance measured in the time0 duplicate samples. Gene fitness is the weighted average of individual strain fitness values (Wetmore et al., 2015). For fitness calculations, insertions in the first and last 10% of coding regions were excluded, with insertions in the remaining 80% of the gene considered “central”. While 91 to 92% of genes in all strains contained centrally mapped insertions, not all genes were used in fitness calculations due to low read or insertion abundance. We focused our analysis on genes with fitness values >1 or <-1, and absolute t-score >4 (Table S6). Across all conditions, we calculated fitness values for 3,705 (*Dda*3937), 3,761 (*Ddia*ME23), and 3,528 (*Ddia*6719) genes, representing 88%, 90%, and 86% of the total genes in each strain respectively.

Principle component analysis showed gene fitness values of the three tuber conditions overlapped (Fig. 1), and so these samples were jointly considered as a single “Tuber” condition for some subsequent analyses.

**Figure 1.**
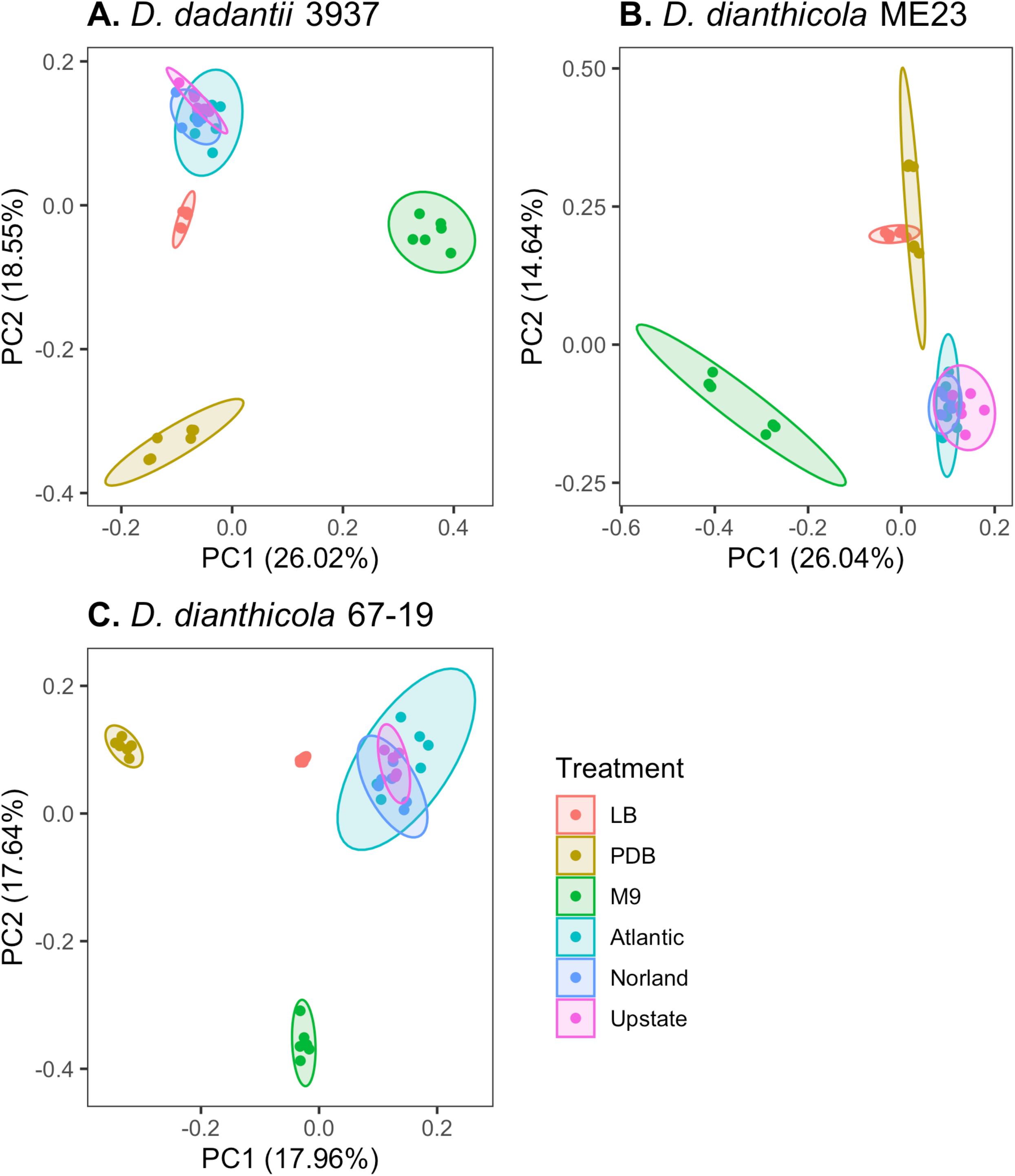
PCA of experimental samples based on fitness values calculated for each library. Available fitness values for each sample: N= 3,705 *D. dadantii* 3937; 3,761 *D. dianthicola* ME23; 3,528 *D. dianthicola* 67-19). Superimposed ellipses are based on a multivariate t-distribution.

### Disruption mutants with fitness defects in rich media

As the libraries were constructed on LB medium, relatively few mutations deleterious in LB were maintained in the populations (Fig. 2). Fig. 3 presents these data split by COG category. Limited mutations in genes categorized as “cell wall/membrane/envelope biogenesis” (*mdoGH*) and “cell cycle control, cell division, chromosome partitioning” (*ftsX*) were present in the mapped populations but generally detrimental in LB for all three strains (Fig. S4). Even in LB, some variation was apparent between strains, such as disruptions in the gene encoding the cell division protein ZapB which decreased competitive fitness in *Dda*3937 but not *Ddia*ME23 or *Ddia*6719 (Fig. S4).

**Figure 2.**
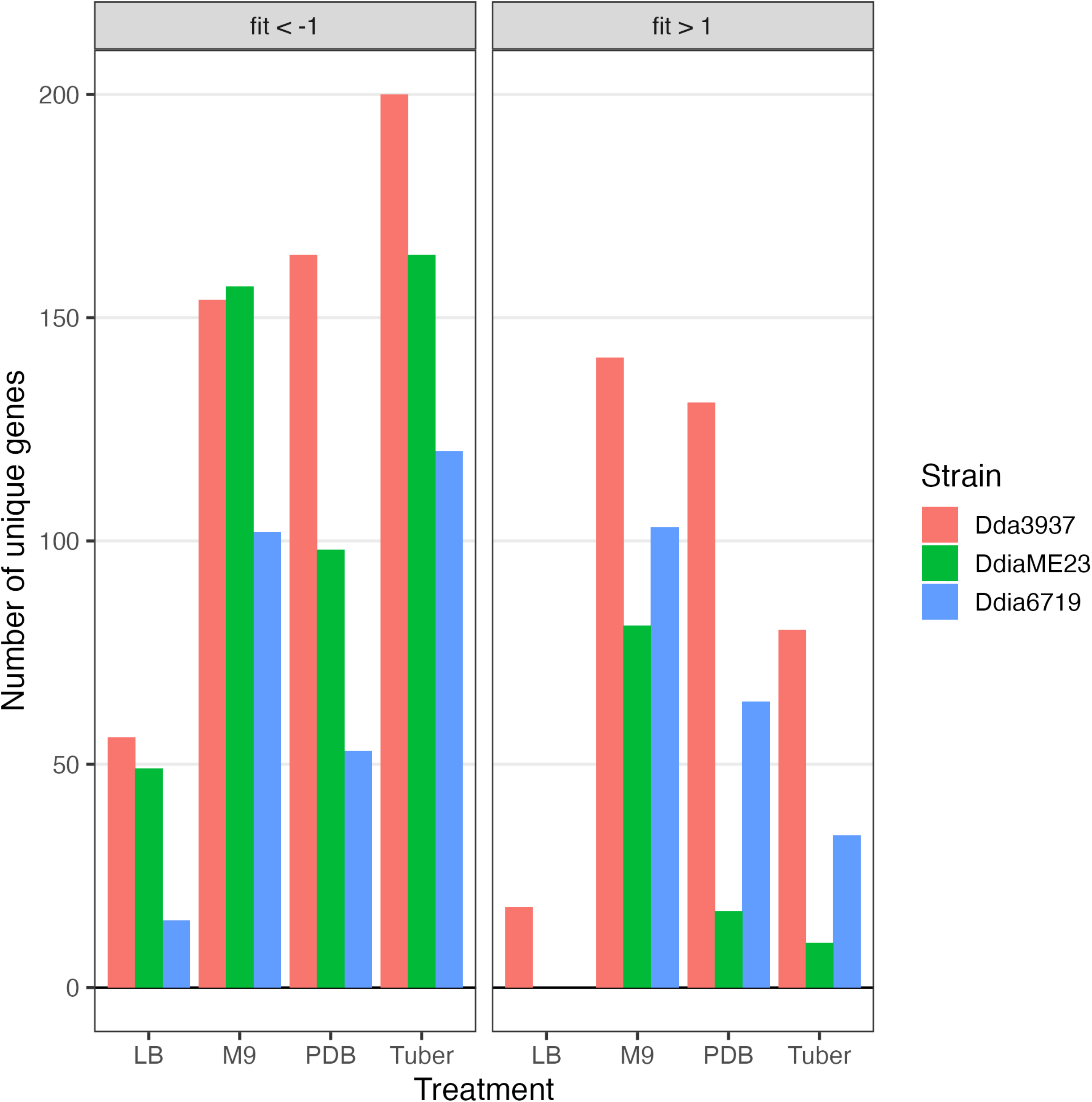
Number of unique genes for each condition with fitness values of <-1 or >1, and absolute t-like test statistic >4 in at least one replicate sample.

**Figure 3.**
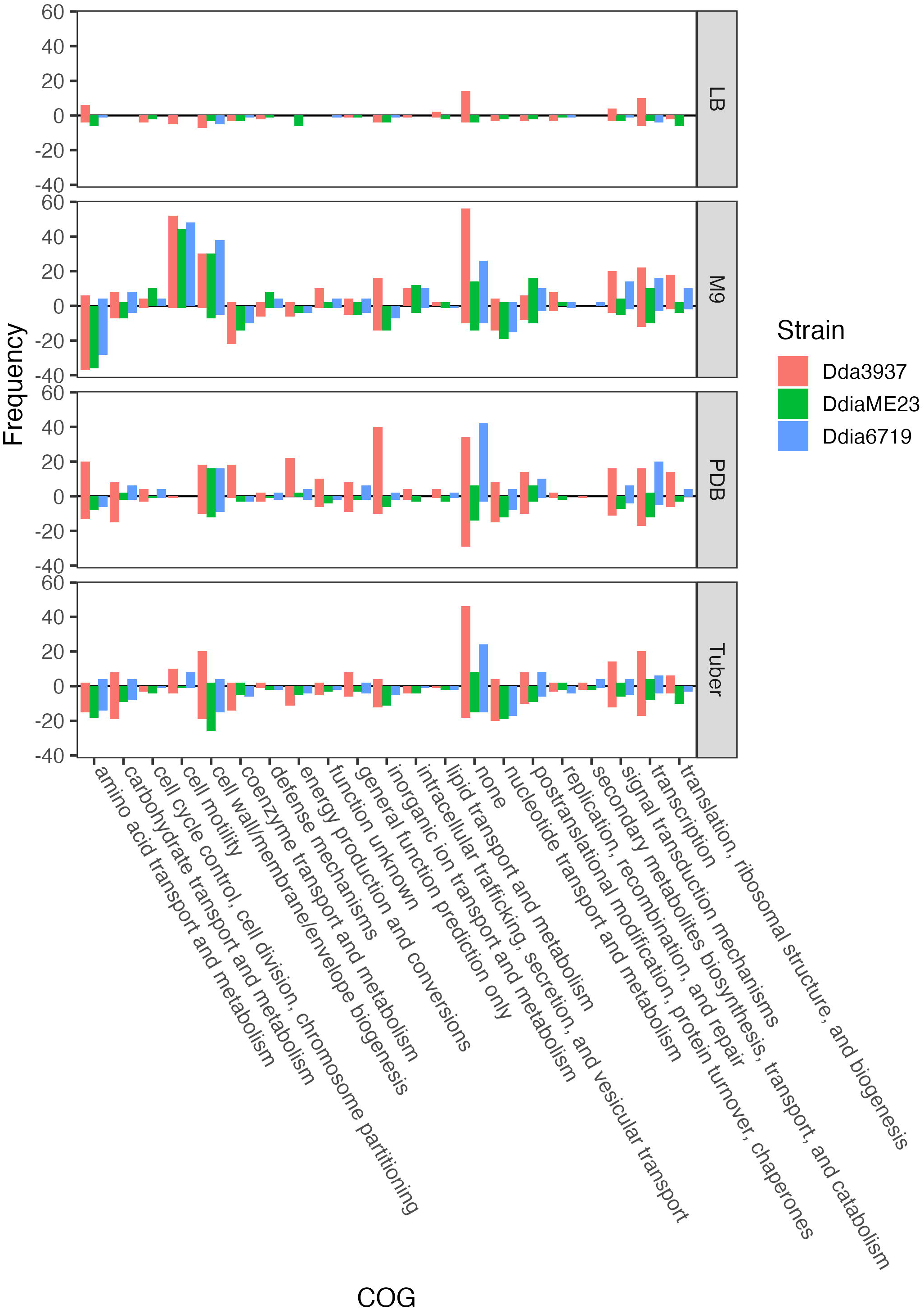
Number of unique genes for each condition with fitness values of <-1 or >1, and absolute t-like test statistic >4 in at least one replicate sample. Genes where fit <-1 are show below the line y=0, while genes where fit >1 are shown above. COG assignments are based on the *D. dadantii* 3937 annotation in the IMG database (Chen et al., 2019), and extrapolated to *D. dianthicola* ME23 and *D. dianthicola* 67-19 based on PyParanoid-generated ortholog groups.

The rich medium PDB provided a very different gene fitness profile than LB. In *Dda*3937 and *Ddia*6719 similar numbers of genes were detrimental (fit >1) as were beneficial (fit <-1) in this condition (Fig. 2). Genes in diverse metabolic categories contributed to competitive fitness, including “amino acid transport and metabolism”, “carbohydrate transport and metabolism”, “cell wall/membrane/envelope biogenesis”, “coenzyme transport and metabolism”, “inorganic ion transport and metabolism”, “nucleotide transport and metabolism”, “signal transduction mechanisms”, “transcription”, and “translation, ribosomal structure, and biogenesis” (Fig. 3). For example, in all three strains oligopeptidase A and the low affinity potassium transporter Kup were specifically important in PDB for growth (Fig. S5). Disruptions in the two-component system RtsAB were specifically beneficial for *Dda*3937 in PDB, as were disruptions in the zinc uptake transcriptional repressor Zur (Fig. S5). Though LB and PDB are both complex rich media, specific available nutrients differed enough to clearly separate the gene fitness profiles for *Dda*3937 and *Ddia*6719, though not for *Ddia*ME23 (Fig. 1).

### Disruption mutants with fitness defects in minimal medium

In the minimal medium M9 containing glycerol as a carbon source, important genes included categories such as “amino acid transport and metabolism”, “carbohydrate transport and metabolism”, “coenzyme transport and metabolism”, and “nucleotide transport and metabolism”. While many amino acids were limiting in both M9 and tuber samples, arginine biosynthetic genes (*argCEFGH*) were uniquely important in M9, suggesting the presence of available arginine in tubers (Fig. 4). Conversely, mutations in many “cell motility” genes had a large positive effect, though this effect was often limited to *Dda*3937 (Fig. 5). This is indicative of the limited energy available in minimal medium, and the high energy cost of motility.

**Figure 4.**
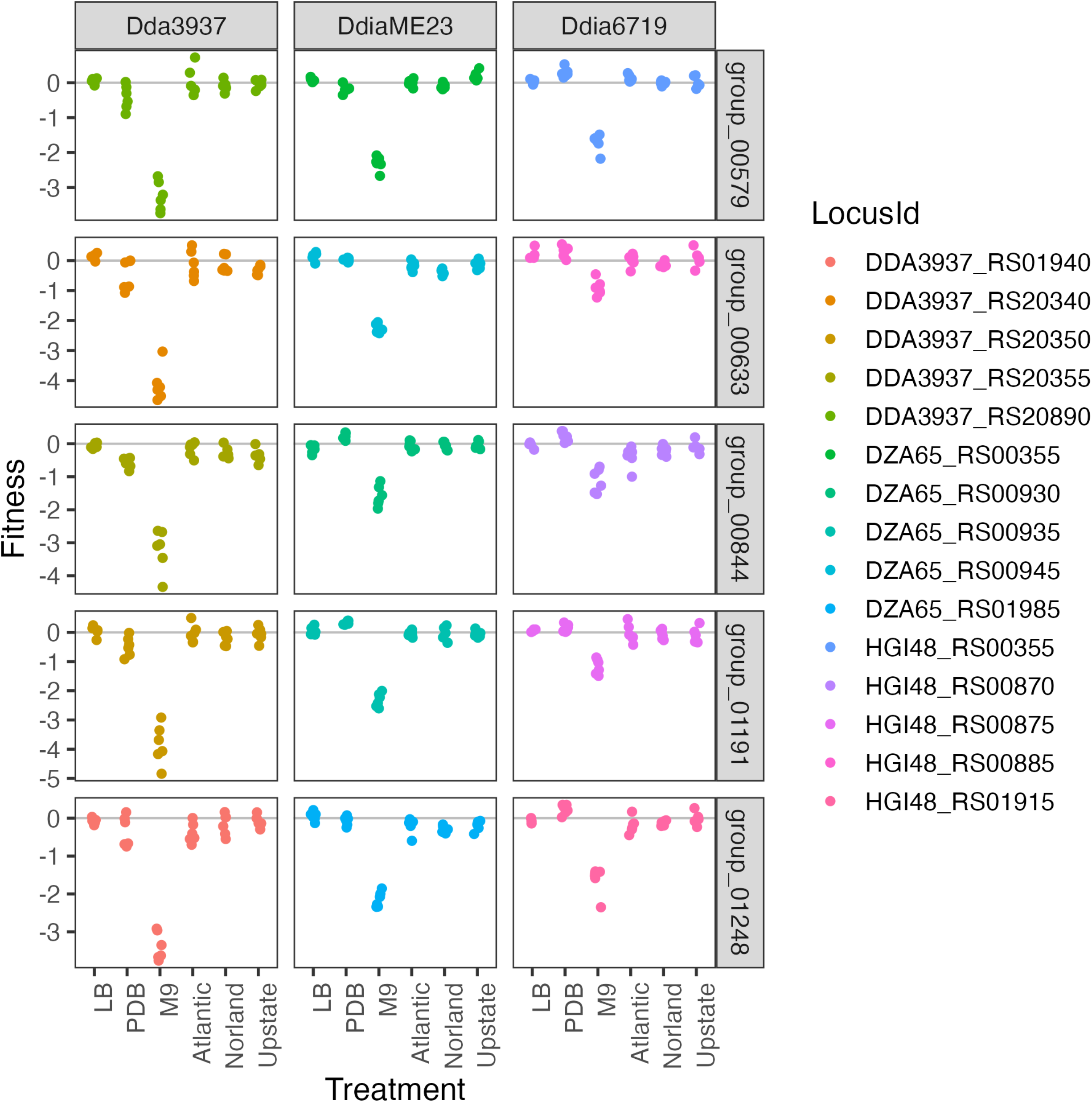
Mutations in arginine biosynthetic genes are detrimental only in M9 minimal medium. Gene fitness values for argininosuccinate synthase ArgG (group 00579), argininosuccinate lyase ArgH (group 00633), acetylornithine deacetylase ArgE (group 00844), N-acetyl-gamma-glutamyl-phosphate reductase ArgC (group 01191), and ornithine carbamoyltransferase ArgF (group 01248).

**Figure 5.**
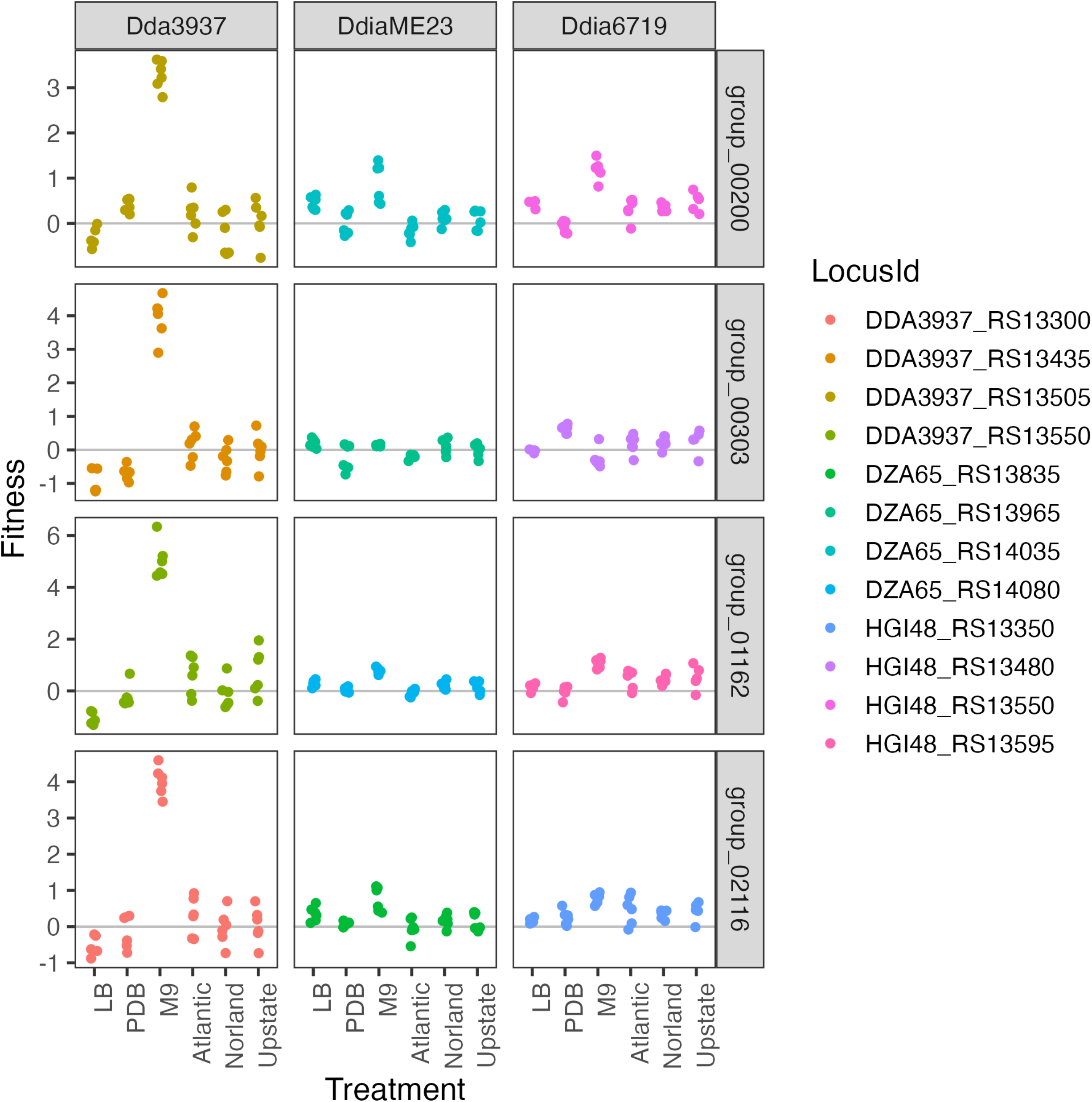
Mutations in flagellar-associated genes increase competitive fitness of *D. dadantii* 3937 in M9 minimal medium. Gene fitness values for flagellar biosynthesis protein FlhA (group 00200), flagellar hook-associated protein FlgK (group 00303), flagellar motor protein MotB (group 001162), and RNA polymerase sigma factor FliA (group 02116.)

### Genes contributing to growth in tubers

To calculate genome-wide gene fitness values in an ecologically- and economically-relevant condition, we inoculated the transposon libraries into tubers of three potato cultivars: “Atlantic”, “Dark Red Norland”, and “Upstate Abundance”. As each transposon library contains over 300,000 unique strains, we inoculated approximately 10^7^ cells into each tuber (10µl of a 10^9^ CFU/ml solution). After two days incubation at high humidity, we recovered cells by streaming for barcode sequencing and calculation of gene fitness values. Many genes involved in amino acid biosynthesis that were important for growth in M9 were also important in tubers (*leuAC, thrC, serB*), highlighting potentially limiting factors for growth during potato soft rot (Fig. 6).

**Figure 6.**
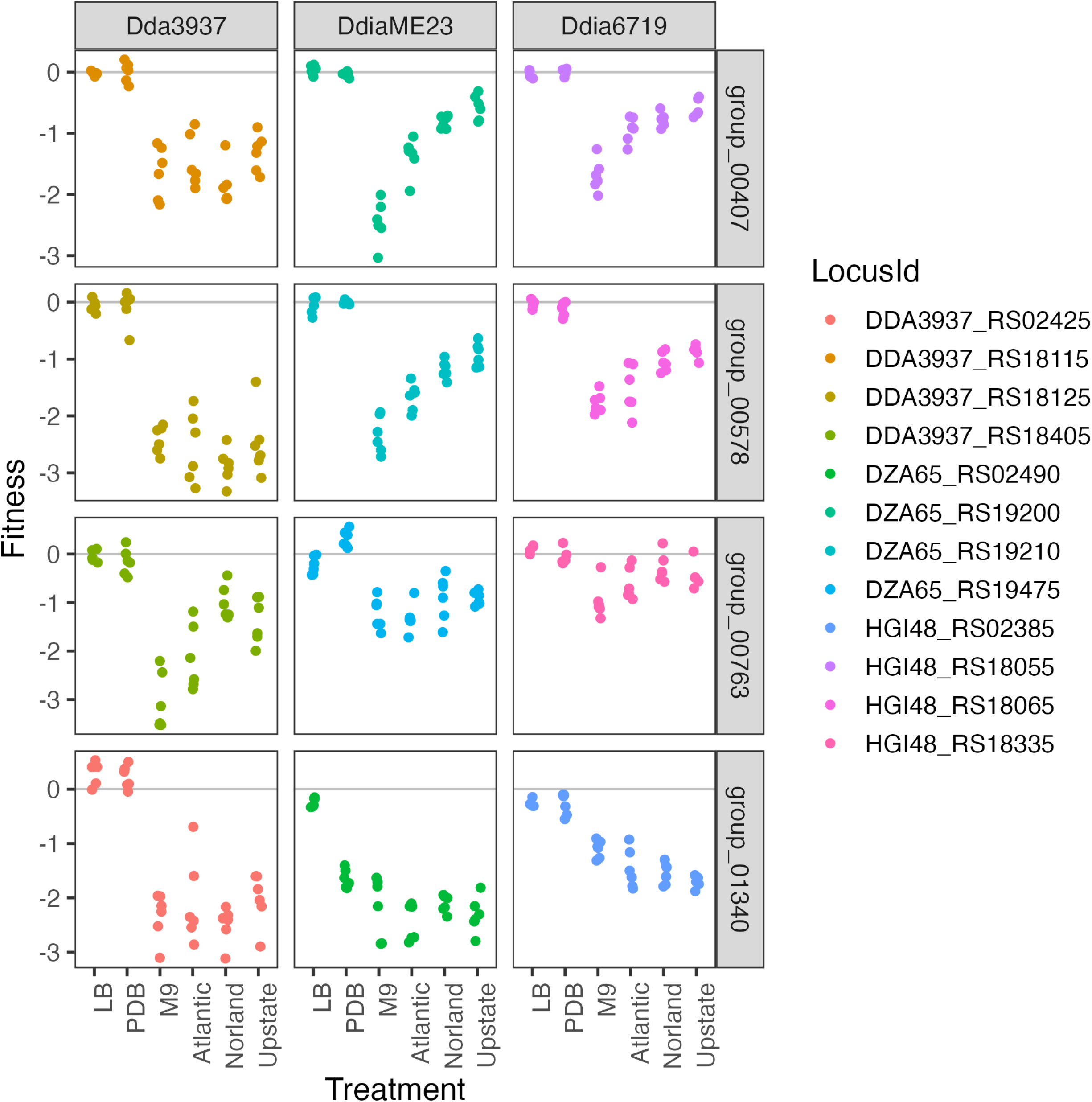
Amino acid biosynthetic genes important in M9 minimal medium as well as growth in potato tubers. Gene fitness values for 2-isopropylmalate synthase LeuA (group 00407), 3-isopropylmalate dehydratase large subunit LeuC (group 00578), threonine synthase ThrC (group 00763), and phosphoserine phosphatase SerB (group 01340).

The pectin degradation protein 2-dehydro-3-deoxy-D-gluconate 5-dehydrogenase KduD was specifically important for growth in tubers (Fig. 7). Interestingly, we identified several putative DNA-binding or helix-turn-helix transcriptional regulators where mutant strains had strain-specific increased fitness in tubers (Fig. 7). Insertions in the *Ddia*6719 helix-turn-helix transcriptional regulator *HGI48_RS01985* increased fitness in tubers, while insertions in the paralog *HGI48_RS02000* had no effect on fitness. There is no ortholog for this gene in *Dda3937*, and the ortholog in *Ddia*ME23 had no disruption phenotype in any condition tested (Fig. 7).

**Figure 7.**
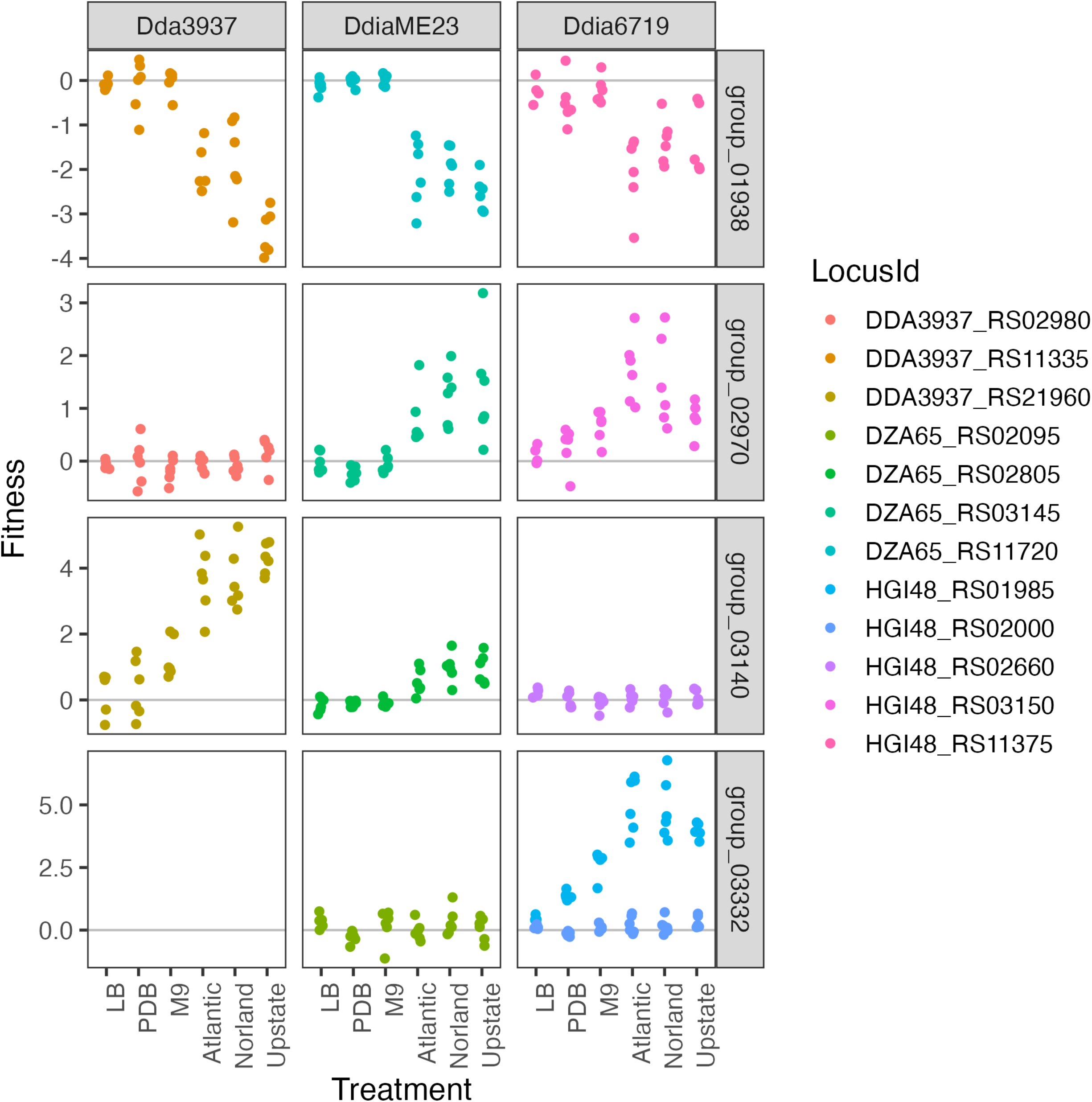
In tubers, pectin degradation contributes to growth, while the role of putative DNA-binding proteins varies among strains. Pectin degradation by 2-dehydro-3-deoxy-D-gluconate 5-dehydrogenase KduD (group 01938) is specifically important for growth in tubers. Gene fitness values shown for three putative DNA binding proteins: ortholog groups 02970, 03140, and 03332. There was no ortholog detected in *D. dadantii* 3937 for group 03332. In *D. dianthicola* 67-19, there are two genes in group 03332. All other orthogroup members are single-copy in each strain.

## Discussion

By measuring genome-wide gene contributions to growth, we identified a comprehensive list of genes in three *Dickeya* strains that contribute to fitness in diverse conditions including potato tubers. Our data generally support previous findings from other groups, including the importance of diverse metabolic capabilities during tuber colonization and the production of pectin degradation proteins (Condemine and Robert□Baudouy, 1991). Royet et al. used TnSeq in the *Dickeya*-chicory pathosystem to identify important genes for growth in leaves (Royet et al., 2019). While there are important differences between the different plant tissue types, our results were very similar overall. Many core metabolic processes were highly important in both minimal medium and tuber samples, such as biosynthesis of many amino acids. However, potato tubers contain higher arginine concentrations than other essential amino acids (Bártová et al., 2015), correlating with the dispensability of the arginine biosynthetic genes (*argCEFGH*) in tubers (Fig. 4). Interestingly, though an *Erwinia amylovora argD* mutant is auxotrophic and nonpathogenic in apple (Ramos et al., 2014), *argD* is present in two apparently redundant copies in *Dda*3937 (DDA3937_RS19450 and DDA3937_RS03635), and mutants in these genes had no phenotype in the conditions tested. Previous studies have shown the important of chemotaxis and motility for early-stage virulence (Jahn et al., 2008), but these traits were dispensable for growth in tubers with the inoculation and sampling methods used here.

While *D. dadantii* and *D. dianthicola* can cause soft rot on potato tubers, variation in some key genes under these conditions suggests species- and strain-level differences in virulence strategies and stress responses. General strategies for environmental growth and host colonization are consistent, such as general metabolic capabilities and stress tolerance. However, gene fitness data suggest variation in gene regulation, such as the helix-turn-helix and other putative DNA-binding proteins. Furthermore, the large fitness increases seen in liquid minimal medium when flagellar genes are disrupted in *Dda*3937 but not *Ddia*ME23 or *Ddia*6719 might indicate weaker control of gene expression, and therefore energy loss and decreased growth when these traits are dispensable or unnecessary. Further characterization of the regulation of these traits is needed.

The scalability of RB-TnSeq, paired with ortholog identification, has proven to be a useful method to directly compare gene fitness between related strains. *Dickeya* species generally have common virulence strategies, primarily the production of plant cell wall degrading enzymes such as pectate lyases (Reverchon et al., 2016). However, genomic and transcriptomic variation at the strain and species level highlights distinctive virulence traits (Raoul des Essarts et al., 2019). This leads to the intriguing possibility that while enzymatic virulence traits are shared across pathogens, there exists strain-specific virulence regulation. This idea has been proposed, but not directly tested, in *D. solani* based on predicted binding sites for transcriptional regulators (Golanowska et al., 2018). In the case of our study, while *Dda*3937 is pathogenic on potato, it was originally isolated from *Saintpaulia ionantha* (Lemattre and Narcy, 1972), suggesting potato infection is simply opportunistic. *Ddia*67-19 was originally isolated from New Guinea impatiens (*Impatiens hawkeri*) (Liu et al., 2020a), but observed symptoms in tubers were similar to those caused by *Ddia*ME23. Based on our fitness data, pan-genome analysis within the *Dickeya* clade might indicate other strains with potentially interesting genome composition for RB-TnSeq analysis.

This study focused on isolated strain growth, to generate a comprehensive dataset of likely essential genes in *D. dadantii* and *D. dianthicola*, and those involved in potato soft rot. Testing other *Dickeya* species, as well as related pathogens such as *Pectobacterium* spp., will more broadly support our understanding of soft rot pathogens. Furthermore, varied additional *in vitro* conditions such as alternative carbon and nitrogen sources can clarify specific metabolic pathways used by these strains for full virulence. In the field, soft rot symptoms can be the result of complex community interactions, with *Dickeya* and *Pectobacterium* co-infections frequently observed (Ge et al., 2021a). It would be interesting to see if the presence of additional community members might change the genes required for full competitive fitness. Though there are no resistant potato cultivars, varieties have been identified with disease tolerance (Lyon, 1989). Tolerance mechanisms being tested include plant cell wall modifications, production of bactericidal proteins and specialized metabolites, and molecules to dysregulate bacterial quorum sensing (Czajkowski et al., 2011). Good hygiene controls at the seed treatment and postharvest stages are also critical for disease mitigation (van der Wolf and De Boer, 2007; Toth et al., 2011). Understanding bacterial virulence strategies will aid in breeding efforts, as well as identify potential bacterial traits that could enable overcoming host tolerance or exacerbating disease at all stages of production.

## Acknowledgements

The authors would like to thank Walter De Jong and the Cornell Potato Breeding Program for providing potato tubers. The authors also thank Adam Deutschbauer for providing the donor *E. coli* library APA752 containing the barcoded *mariner* vector and Morgan Price for assisting with the FEBA pipeline and data upload to the Fitness Browser. Sequencing was performed by the Biotechnology Resource Center (BRC) Genomics Facility at the Cornell Institute of Biotechnology. Mention of trade names or commercial products in this publication is solely for the purpose of providing specific information and does not imply recommendation or endorsement by the U.S. Department of Agriculture. USDA is an equal opportunity provider and employer.

## Notes

https://fit.genomics.lbl.gov/

https://github.com/tylerhelmann/dickeya-barseq-2021

### Competing Interest Statement

The authors have declared no competing interest.

## References

Adeolu, M., Alnajar, S., Naushad, S., and Gupta, R. S. (2016). Genome-based phylogeny and taxonomy of the ‘Enterobacteriales’: Proposal for Enterobacterales ord. nov. divided into the families Enterobacteriaceae, Erwiniaceae fam. nov., Pectobacteriaceae fam. nov., Yersiniaceae fam. nov., <. Int. J. Syst. Evol. Microbiol. 66, 5575–5599. doi:10.1099/ijsem.0.001485.

Adriaenssens, E. M., van Vaerenbergh, J., Vandenheuvel, D., Dunon, V., Ceyssens, P. J., de Proft, M., et al. (2012). T4-related bacteriophage LIMEstone isolates for the control of soft rot on potato caused by “Dickeya solani.” PLoS One 7, e33227. doi:10.1371/journal.pone.0033227.

Bártová, V., Bárta, J., Brabcová, A., Zdráhal, Z., and Horáčková, V. (2015). Amino acid composition and nutritional value of four cultivated South American potato species. J. Food Compos. Anal. 40, 78–85. doi:10.1016/j.jfca.2014.12.006.

Bertani, G. (1951). Studies on lysogenesis. I. The mode of phage liberation by lysogenic Escherichia coli. J. Bacteriol. 62, 293–300. doi:10.1128/JB.62.3.293-300.1951.

Chen, I. M. A., Chu, K., Palaniappan, K., Pillay, M., Ratner, A., Huang, J., et al. (2019). IMG/M v.5.0: an integrated data management and comparative analysis system for microbial genomes and microbiomes. Nucleic Acids Res. 47, D666–D677. doi:10.1093/nar/gky901.

Chung, Y. S., Goeser, N. J., Cai, X., and Jansky, S. (2013). The effect of long term storage on bacterial soft rot resistance in potato. Am. J. Potato Res. 90, 351–356. doi:10.1007/s12230-013-9311-6.

Cole, B. J., Feltcher, M. E., Waters, R. J., Wetmore, K. M., Mucyn, T. S., Ryan, E. M., et al. (2017). Genome-wide identification of bacterial plant colonization genes. PLoS Biol. 15, 1– 24. doi:10.1371/journal.pbio.2002860.

Condemine, G., and Robert□Baudouy, J. (1991). Analysis of an Erwinia chrysanthemi gene cluster involved in pectin degradation. Mol. Microbiol. 5, 2191–2202. doi:10.1111/j.1365-2958.1991.tb02149.x.

Czajkowski, R., Pérombelon, M. C. M., Van Veen, J. A., and Van der Wolf, J. M. (2011). Control of blackleg and tuber soft rot of potato caused by Pectobacterium and Dickeya species: A review. Plant Pathol. 60, 999–1013. doi:10.1111/j.1365-3059.2011.02470.x.

Czajkowski, R., Smolarska, A., and Ozymko, Z. (2017). The viability of lytic bacteriophage ΦD5 in potato-associated environments and its effect on Dickeya solani in potato (Solanum tuberosum L.) plants. PLoS One 12, 1–17. doi:10.1371/journal.pone.0183200.

Ge, T., Ekbataniamiri, F., Johnson, S. B., Larkin, R. P., and Hao, J. (2021a). Interaction between Dickeya dianthicola and Pectobacterium parmentieri in potato infection under field conditions. Microorganisms 9, 1–10. doi:10.3390/microorganisms9020316.

Ge, T., Jiang, H., Tan, E. H., Johnson, S. B., Larkin, R., Charkowski, A. O., et al. (2021b). Pangenomic analysis of Dickeya dianthicola strains related to the outbreak of blackleg and soft rot of potato in USA. Plant Dis., 1–31. doi:10.1094/PDIS-03-21-0587-RE.

Georgoulis, S., Shalvarjian, K. E., Helmann, T. C., Hamilton, C. D., Carlson, H. K., Deutschbauer, A. M., et al. (2020). Genome-wide identification of tomato xylem sap fitness factors for Ralstonia pseudosolanacearum and Ralstonia syzygii. bioRxiv, 1–45. doi:https://doi.org/10.1101/2020.08.31.276741.

Golanowska, M., Potrykus, M., Motyka-Pomagruk, A., Kabza, M., Bacci, G., Galardini, M., et al. (2018). Comparison of highly and weakly virulent Dickeya solani strains, with a view on the pangenome and panregulon of this species. Front. Microbiol. 9, 1–19. doi:10.3389/fmicb.2018.01940.

Helmann, T. C., Deutschbauer, A. M., and Lindow, S. E. (2019). Genome-wide identification of Pseudomonas syringae genes required for fitness during colonization of the leaf surface and apoplast. Proc. Natl. Acad. Sci. 116, 18900–18910. doi:10.1073/pnas.1908858116.

Jahn, C. E., Willis, D. K., and Charkowski, A. O. (2008). The flagellar sigma factor FliA is required for Dickeya dadantii virulence. Mol. Plant-Microbe Interact. 21, 1431–1442. doi:10.1094/MPMI-21-11-1431.

Jiang, X., Zghidi-Abouzid, O., Oger-Desfeux, C., Hommais, F., Greliche, N., Muskhelishvili, G., et al. (2016). Global transcriptional response of Dickeya dadantii to environmental stimuli relevant to the plant infection. Environ. Microbiol. 18, 3651–3672. doi:10.1111/1462-2920.13267.

Lee, D. H., Lim, J. A., Lee, J., Roh, E., Jung, K., Choi, M., et al. (2013). Characterization of genes required for the pathogenicity of Pectobacterium carotovorum subsp. carotovorum Pcc21 in Chinese cabbage. Microbiol. (United Kingdom) 159, 1487–1496. doi:10.1099/mic.0.067280-0.

Lemattre, M., and Narcy, J. P. (1972). Une affection bacterienne nouvelle du Saintpaulia due a Erwinia chrysanthemi. C. R. Acad. Sci 58, 227–231.

Liu, H., Shiver, A. L., Price, M. N., Carlson, H. K., Trotter, V. V., Chen, Y., et al. (2021). Functional genetics of human gut commensal Bacteroides thetaiotaomicron reveals metabolic requirements for growth across environments. Cell Rep. 34, 108789. doi:10.1016/j.celrep.2021.108789.

Liu, Y., Helmann, T., Stodghill, P., and Filiatrault, M. (2020a). Complete genome sequence resource for the necrotrophic plant-pathogenic bacterium Dickeya dianthicola 67-19 isolated from New Guinea Impatiens. Plant Dis., PDIS-09-20-1968-A. doi:10.1094/PDIS-09-20-1968-A.

Liu, Y., Vasiu, S., Daughtrey, M. L., and Filiatrault, M. (2020b). First Report of Dickeya dianthicola causing blackleg on New Guinea Impatiens (Impatiens hawkeri) in New York State, USA. Plant Dis. doi:10.1094/pdis-09-20-2020-pdn.

Lyon, G. D. (1989). The biochemical basis of resistance of potatoes to soft rot Erwinia spp.—a review. Plant Pathol. 38, 313–339. doi:10.1111/j.1365-3059.1989.tb02152.x.

M9 minimal medium (standard) (2010). Cold Spring Harb. Protoc. 2010, pdb.rec12295. doi:10.1101/pdb.rec12295.

Ma, X., Perna, N. T., Glasner, J. D., Hao, J., Johnson, S., Nasaruddin, A. S., et al. (2019). Complete genome sequence of Dickeya dianthicola ME23, a pathogen causing blackleg and soft rot diseases of potato. Microbiol. Resour. Announc. 8, 14–15. doi:10.1128/mra.01526-18.

Melnyk, R. A., Hossain, S. S., and Haney, C. H. (2019). Convergent gain and loss of genomic islands drives lifestyle changes in plant-associated Pseudomonas. ISME J. 13, 1575–1588. doi:10.1101/345488.

Motyka-Pomagruk, A., Zoledowska, S., Misztak, A. E., Sledz, W., Mengoni, A., and Lojkowska, E. (2020). Comparative genomics and pangenome-oriented studies reveal high homogeneity of the agronomically relevant enterobacterial plant pathogen Dickeya solani. BMC Genomics 21, 1–18. doi:10.1186/s12864-020-06863-w.

Motyka, A., Zoledowska, S., Sledz, W., and Lojkowska, E. (2017). Molecular methods as tools to control plant diseases caused by Dickeya and Pectobacterium spp: A minireview. N. Biotechnol. 39, 181–189. doi:10.1016/j.nbt.2017.08.010.

Parkinson, N., DeVos, P., Pirhonen, M., and Elphinstone, J. (2014). Dickeya aquatica sp. nov., isolated from waterways. Int. J. Syst. Evol. Microbiol. 64, 2264–2266. doi:10.1099/ijs.0.058693-0.

Price, M. N., Wetmore, K. M., Waters, R. J., Callaghan, M., Ray, J., Liu, H., et al. (2018). Mutant phenotypes for thousands of bacterial genes of unknown function. Nature 557, 503– 509. doi:10.1038/s41586-018-0124-0.

R Core Team (2017). R: A language and environment for statistical computing. Vienna, Austria: R Foundation for Statistical Computing doi:10.1007/978-3-540-74686-7.

Ramos, L. S., Lehman, B. L., Peter, K. A., and McNellis, T. W. (2014). Mutation of the Erwinia amylovora argD gene causes arginine auxotrophy, nonpathogenicity in apples, and reduced virulence in pears. Appl. Environ. Microbiol. 80, 6739–6749. doi:10.1128/AEM.02404-14.

Raoul des Essarts, Y., Pédron, J., Blin, P., Van Dijk, E., Faure, D., and Van Gijsegem, F. (2019). Common and distinctive adaptive traits expressed in Dickeya dianthicola and Dickeya solani pathogens when exploiting potato plant host. Environ. Microbiol. 21, 1004–1018. doi:10.1111/1462-2920.14519.

Reverchon, S., Muskhelisvili, G., and Nasser, W. (2016). Virulence Program of a Bacterial Plant Pathogen: The Dickeya Model. Elsevier Inc. doi:10.1016/bs.pmbts.2016.05.005.

Royet, K., Parisot, N., Rodrigue, A., Gueguen, E., and Condemine, G. (2019). Identification by Tn-seq of Dickeya dadantii genes required for survival in chicory plants. Mol. Plant Pathol. 20, 287–306. doi:10.1111/mpp.12754.

Rubin, B. E., Diamond, S., Cress, B. F., Crits-Christoph, A., He, C., Xu, M., et al. (2020). Targeted genome editing of bacteria within microbial communities. bioRxiv, 1–49. doi:10.1101/2020.07.17.209189.

Samson, R., Legendre, J. B., Christen, R., Fischer-Le Saux, M., Achouak, W., and Gardan, L. (2005). Transfer of Pectobacterium chrysanthemi (Burkholder et al. 1953) Brenner et al. 1973 and Brenneria paradisiaca to the genus Dickeya gen. nov. as Dickeya chrysanthemi comb. nov. and Dickeya paradisiaca comb. nov. and deli. Int. J. Syst. Evol. Microbiol. 55, 1415–1427. doi:10.1099/ijs.0.02791-0.

Tian, Y., Zhao, Y., Yuan, X., Yi, J., Fan, J., Xu, Z., et al. (2016). Dickeya fangzhongdai sp. nov., a plant-pathogenic bacterium isolated from pear trees (Pyrus pyrifolia). Int. J. Syst. Evol. Microbiol. 66, 2831–2835. doi:10.1099/ijsem.0.001060.

Toth, I. K., van der Wolf, J. M., Saddler, G., Lojkowska, E., Hélias, V., Pirhonen, M., et al. (2011). Dickeya species: An emerging problem for potato production in Europe. Plant Pathol. 60, 385–399. doi:10.1111/j.1365-3059.2011.02427.x.

van der Wolf, J. M., and De Boer, S. H. (2007). Bacterial pathogens of potato. Elsevier B.V. doi:10.1016/B978-044451018-1/50069-5.

Van Gijsegem, F., Toth, I. K., and van der Wolf, J. M. (2021). “Outlook-Challenges and perspectives for management of diseases caused by Pectobacterium and Dickeya species,” in Plant Diseases Caused by Dickeya and Pectobacterium Species, eds. F. Van Gijsegem, J. M. van der Wolf, and I. K. Toth (Cham: Springer International Publishing), 283–289. doi:10.1007/978-3-030-61459-1_9.

van Opijnen, T., Bodi, K. L., and Camilli, A. (2009). Tn-seq: high-throughput parallel sequencing for fitness and genetic interaction studies in microorganisms. Nat. Methods 6, 767–772. doi:10.1038/nmeth.1377.

Wetmore, K. M., Price, M. N., Waters, R. J., Lamson, J. S., He, J., Hoover, C. A., et al. (2015). Rapid quantification of mutant fitness in diverse bacteria by sequencing randomly bar-coded transposons. MBio 6, 1–15. doi:10.1128/mBio.00306-15.

Wickham, H. (2016). ggplot2: Elegant Graphics for Data Analysis. Springer-Verlag New York Available at: https://ggplot2.tidyverse.org.

Wolf, J. M. Van Der, Nijhuis, E. H., Kowalewska, M. J., Saddler, G. S., Parkinson, N., Elphinstone, J. G., et al. (2014). Dickeya solani sp. nov., a pectinolytic plant-pathogenic bacterium isolated from potato (Solanum tuberosum). Int. J. Syst. Evol. Microbiol. 64, 768– 774. doi:10.1099/ijs.0.052944-0.

Zhang, Y., Fan, Q., and Loria, R. (2016). A re-evaluation of the taxonomy of phytopathogenic genera Dickeya and Pectobacterium using whole-genome sequencing data. Syst. Appl. Microbiol. 39, 252–259. doi:10.1016/j.syapm.2016.04.001.

