## Supplementary Information for "Genome-wide identification of genes important for growth of *Dickeya dadantii* and *D. dianthicola* in potato (*Solanum tuberosum*) tubers"

**Supplemental Information**

This file includes:

- Tables S1 to S6
- Figures S1 to S5
- SI References

Supplemental tables S2, S3, S4, and S6 are attached as separate spreadsheets.

**Supplementary Table 1.** Strains used in this study.

| Strain | Description | Source |
|---|---|---|
| *D. dadantii* 3937 | Wild-type strain | (Lemattre and Narcy 1972; Samson et al. 2005) |
| *D. dianthicola* ME23 | Wild-type strain | (Ma et al. 2019) |
| *D. dianthicola* 67-19 | Wild-type strain | (Liu et al. 2020) |
| *E. coli* WM3064 | Strain APA752; barcoded *mariner* transposon vector ($Kan^R$) in *E. coli* conjugation strain | (Wetmore et al. 2015) |
| *D. dadantii* 3937 | Whole genome barcoded *mariner* transposon library ($Kan^R$) | This work |
| *D. dianthicola* ME23 | Whole genome barcoded *mariner* transposon library ($Kan^R$) | This work |
| *D. dianthicola* 67-19 | Whole genome barcoded *mariner* transposon library ($Kan^R$) | This work |

**Supplementary Table 2.** Individual Sequence Read Archive (SRA) sample accession numbers for BioProject accession PRJNA692477. Reads used for library mapping include the strain name in the file name. BarSeq reads are named by the sample index "TCH#".

**Supplementary Table 3.** Gene homolog groups for *Dda*3937, *Ddia*ME23, and *Ddia*6719, calculated using the PyParanoid pipeline (Melnyk et al. 2019). The standard pipeline output includes group numbers and peptide sequence identifiers. This table has gene locus names added as well as COG categories based on the genome sequence annotation of *Dda*3937 from the IMG database (Chen et al. 2019). Gene descriptions for each orthogroup are included in a separate sheet.

**Supplementary Table 4.** Predictions of essential or nearly essential genes in LB, based on the TnSeq data. Protein coding genes were considered essential or nearly essential for growth in LB if normalized read density (reads/nucleotides across the entire gene) and normalized insertion density (sites/nucleotides in the central 10 to 90% of the gene) were below 0.2 (Price et al. 2018). Genes shorter than 175 bp (*Dda*3937) or 150 bp (*Ddia*ME23 and *Ddia*6719) were excluded from this analysis. Functional category annotations are COG assignments for *Dda*3937 genes in the IMG database (Chen et al. 2019), matched to PyParanoid-determined *Ddia*ME23 and *Ddia*6719 orthologs.

**Supplementary Table 5.** Number of genes within each functional category predicted to be essential or nearly essential in LB, based on the TnSeq data. Protein coding genes were considered essential or nearly essential for growth in LB if normalized read density (reads/nucleotides across the entire gene) and normalized insertion density (sites/nucleotides in the central 10 to 90% of the gene) were below 0.2 (Price et al. 2018). Genes shorter than 175 bp (*Dda*3937) or 150 bp (*Ddia*ME23 and *Ddia*6719) were excluded from this analysis. Functional category annotations are COG assignments for *Dda*3937 genes in the IMG database (Chen et al. 2019), matched to PyParanoid-determined *Ddia*ME23 and *Ddia*6719 orthologs.

| **COG** | ***Dda*3937** | ***Ddia*ME23** | ***Ddia*6719** |
|---|---|---|---|
| Translation, ribosomal structure, and biogenesis | 87 | 102 | 102 |
| Cell wall/membrane/envelope biogenesis | 54 | 53 | 51 |
| Coenzyme transport and metabolism | 27 | 37 | 35 |
| Energy production and conversions | 26 | 26 | 24 |
| Replication, recombination, and repair | 25 | 29 | 30 |
| Lipid transport and metabolism | 23 | 23 | 23 |
| Cell cycle control, cell division, chromosome partitioning | 15 | 13 | 14 |
| Nucleotide transport and metabolism | 15 | 13 | 14 |
| None | 14 | 15 | 19 |
| Transcription | 14 | 15 | 15 |
| Intracellular trafficking, secretion, and vesicular transport | 12 | 9 | 10 |
| Posttranslational modification, protein turnover, chaperones | 11 | 15 | 13 |
| Amino acid transport and metabolism | 10 | 18 | 15 |
| Carbohydrate transport and metabolism | 10 | 13 | 12 |
| Inorganic ion transport and metabolism | 6 | 8 | 6 |
| General function prediction only | 4 | 2 | 3 |
| Secondary metabolites biosynthesis, transport, and catabolism | 3 | 2 | 2 |
| Signal transduction mechanisms | 3 | 5 | 5 |
| Function unknown | 2 | 2 | 5 |
| Defense mechanisms | 1 | 1 | 1 |
| RNA processing and modification | 1 | 1 | 1 |
| Cell motility | 0 | 0 | 0 |
| Extracellular structures | 0 | 0 | 0 |
| Mobilome: prophages, transposons | 0 | 0 | 0 |
| (No COG and/or orthogroup assignment) | 11 | 24 | 26 |
| **Total** | **374** | **426** | **426** |

**Supplementary Table 6.** Fitness values and t-scores for all genes where fit >1 or <-1 and absolute t-score >4. Column values include the locus name ("locusId"), the gene product description ("desc"), the experimental sample ("name"), a shortened description where samples with the same name are replicates ("short"), the normalized fitness value ("lrn"), and the t-like test statistic that gives an estimate of the significance of the gene fitness value ("t").

**Supplementary Figure 1.** Distribution of mapped strains for barcoded *mariner* transposon insertion mutant libraries in *D. dadantii* 3937 (Ddi3937), *D. dianthicola* ME23 (DdiaME23), and *D. dianthicola* 67-19 (Ddia6719).

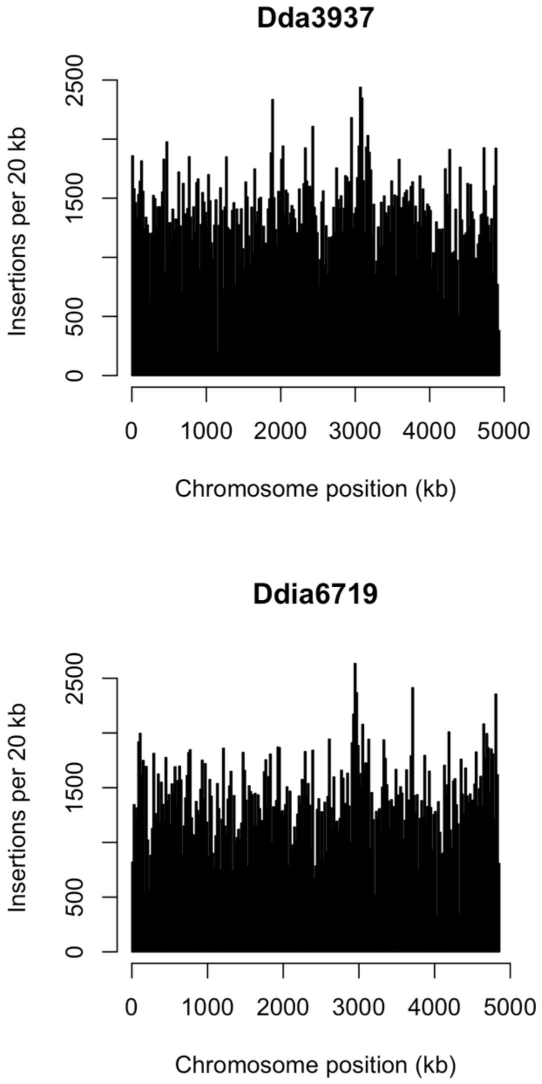

**Supplementary Figure 2.** Most predicted essential genes are common among all three strains. Overlap was calculated based on PyParanoid group numbers for each predicted essential gene. Due to the insertion density of each library, to predict a gene as essential size cutoffs of >175 bp (*Dda*3937) and >150 bp (*Ddia*ME23 and *Ddia*6719) were used. Genes with no ortholog group assignments are not included here: N= 11 (*Dda*3937), 16 (*Ddia*ME23), and 19 (*Ddia*6719).

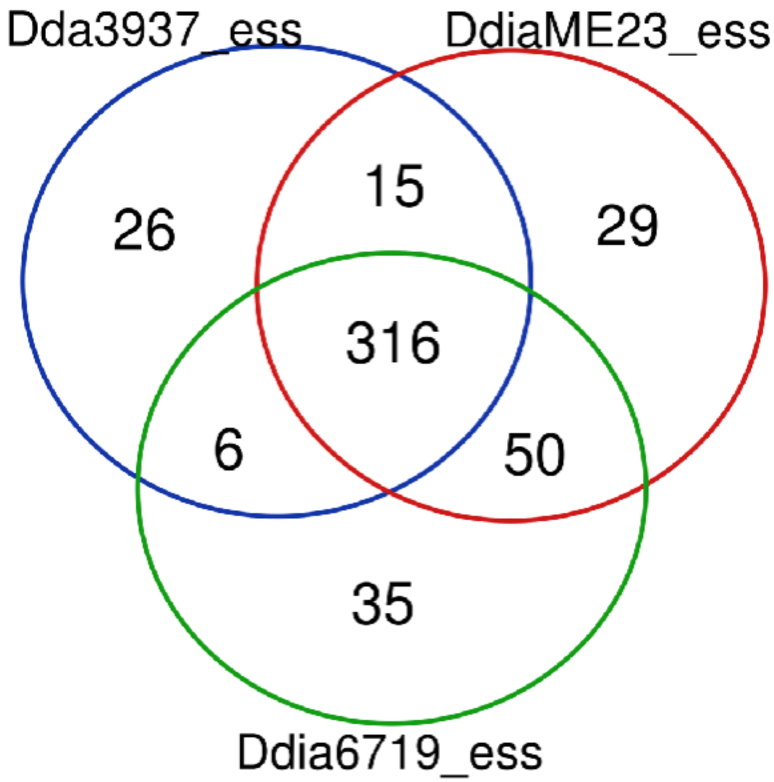

**Supplementary Figure 3.** *In vitro* growth of wild-type *D. dadantii* 3937, *D. dianthicola* ME23, and *D. dianthicola* 67-19 in LB, Potato Dextrose Broth (PDB), and M9 minimal medium containing 0.4% glycerol. (**A**) Growth curve measure absorbance at 600 nm of each strain showing 6 replicate samples per media type. (**B**) Doubling time summary statistics calculated using the R package growthcurver v0.3.1 (Sprouffske and Wagner 2016).

A.

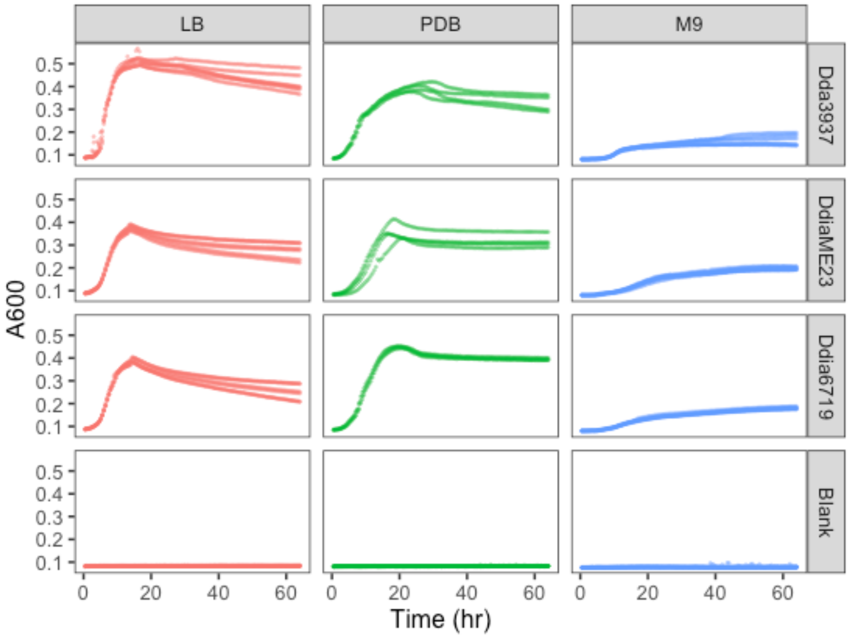


B.

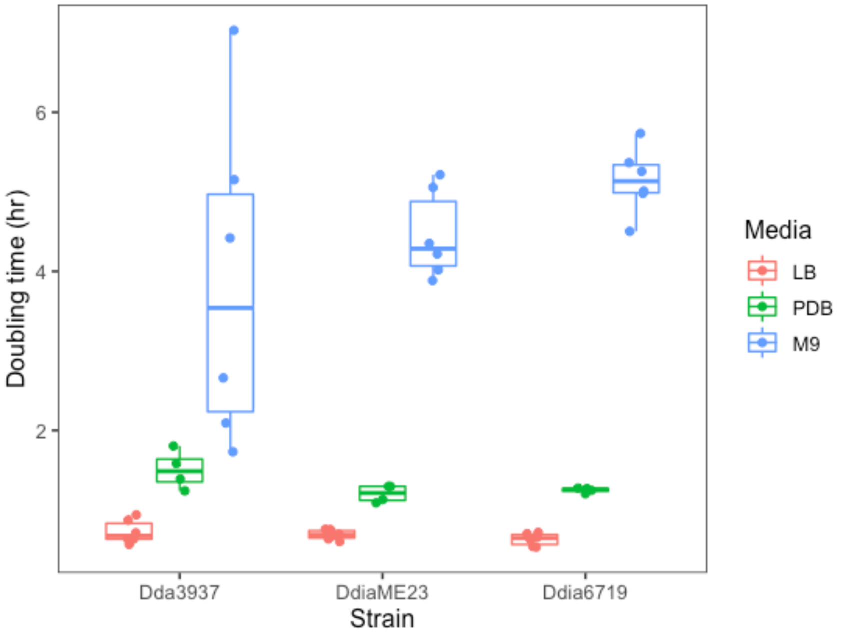

**Supplementary Figure 4.** Gene fitness values for glucans biosynthesis proteins MdoGH (groups 00352 and 00097), cell division protein FtsX (group 01645), and cell division protein ZapB (group 03211).

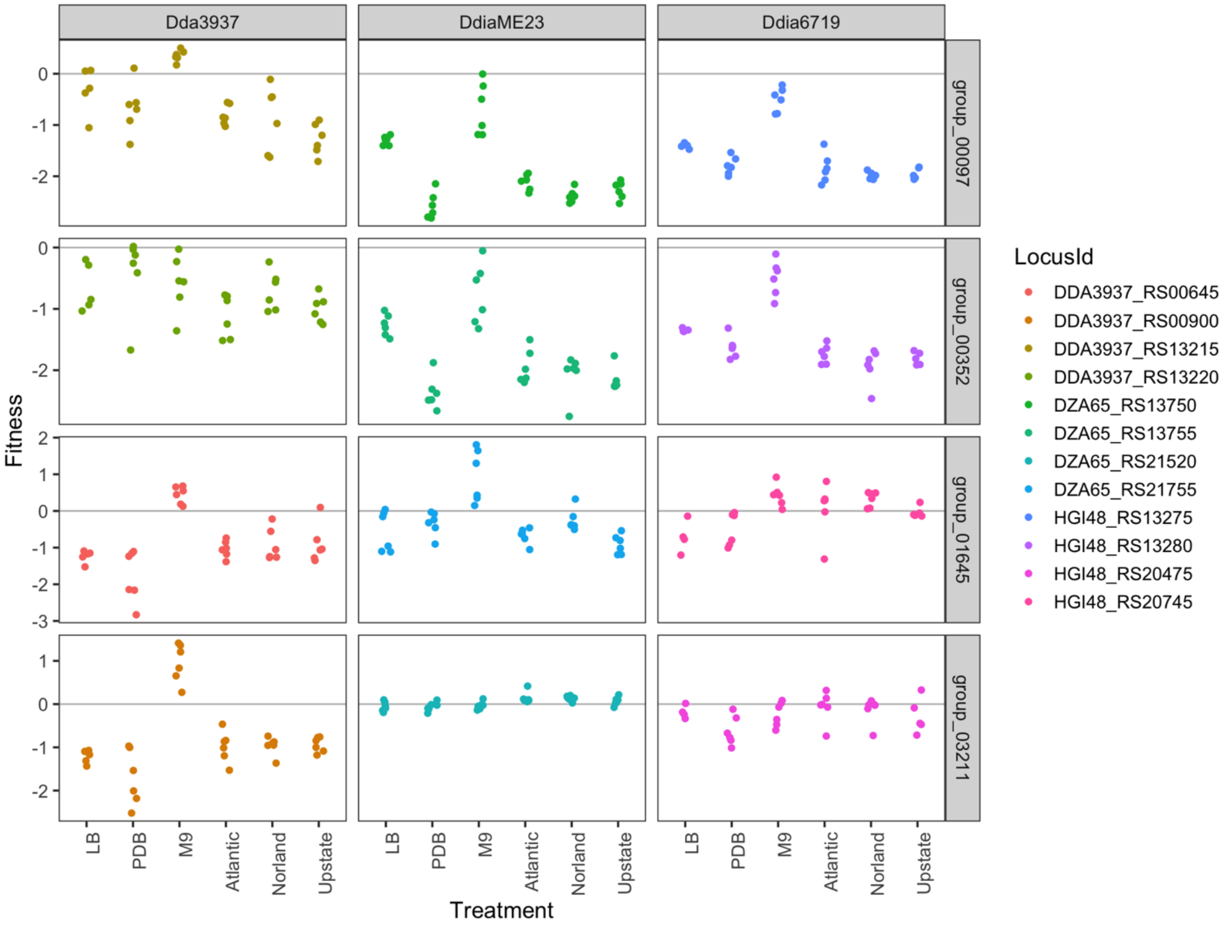

**Supplementary Figure 5.** Gene fitness values for oligopeptidase A (group 00188), the low affinity potassium transporter Kup (group 00240), the two-component system RtsAB (groups 02048 and 00671), and the zinc uptake transcriptional repressor Zur (group 02565).

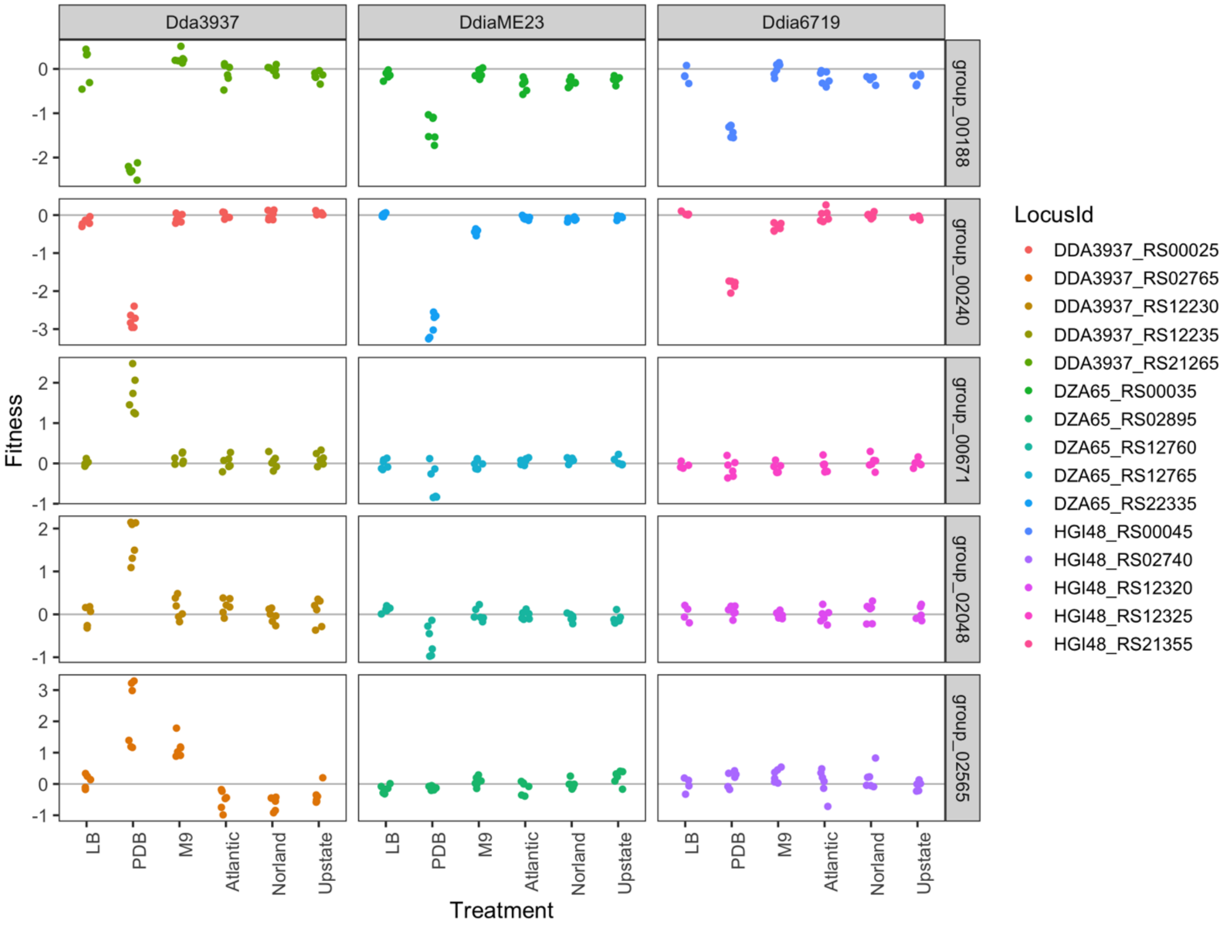

### References


Chen, I. M. A., Chu, K., Palaniappan, K., Pillay, M., Ratner, A., Huang, J., Huntemann, M., Varghese, N., White, J. R., Seshadri, R., Smirnova, T., Kirton, E., Jungbluth, S. P., Woyke, T., Eloe-Fadrosh, E. A., Ivanova, N. N., and Kyrpides, N. C. 2019. IMG/M v.5.0: an integrated data management and comparative analysis system for microbial genomes and microbiomes. Nucleic Acids Res. 47:D666–D677

Lemattre, M., and Narcy, J. P. 1972. Une affection bacterienne nouvelle du Saintpaulia due a *Erwinia chrysanthemi*. C. R. Acad. Sci. 58:227–231

Liu, Y., Helmann, T., Stodghill, P., and Filiatrault, M. 2020. Complete genome sequence resource for the necrotrophic plant-pathogenic bacterium *Dickeya dianthicola* 67-19 isolated from New Guinea Impatiens. Plant Dis. :PDIS-09-20-1968-A

Ma, X., Perna, N. T., Glasner, J. D., Hao, J., Johnson, S., Nasaruddin, A. S., Charkowski, A. O., Wu, S., Fei, Z., Perry, K. L., Stodghill, P., and Swingle, B. 2019. Complete genome sequence of *Dickeya dianthicola* ME23, a pathogen causing blackleg and soft rot diseases of potato. Microbiol. Resour. Announc. 8:14–15

Melnyk, R. A., Hossain, S. S., and Haney, C. H. 2019. Convergent gain and loss of genomic islands drives lifestyle changes in plant-associated *Pseudomonas*. ISME J. 13:1575–1588

Price, M. N., Wetmore, K. M., Waters, R. J., Callaghan, M., Ray, J., Liu, H., Kuehl, J. V., Melnyk, R. A., Lamson, J. S., Suh, Y., Carlson, H. K., Esquivel, Z., Sadeeshkumar, H., Chakraborty, R., Zane, G. M., Rubin, B. E., Wall, J. D., Visel, A., Bristow, J., Blow, M. J., Arkin, A. P., and Deutschbauer, A. M. 2018. Mutant phenotypes for thousands of bacterial genes of unknown function. Nature. 557:503–509

Samson, R., Legendre, J. B., Christen, R., Fischer-Le Saux, M., Achouak, W., and Gardan, L. 2005. Transfer of *Pectobacterium chrysanthemi* (Burkholder et al. 1953) Brenner et al. 1973 and *Brenneria paradisiaca* to the genus *Dickeya* gen. nov. as *Dickeya chrysanthemi* comb. nov. and *Dickeya paradisiaca* comb. nov. and deli. Int. J. Syst. Evol. Microbiol. 55:1415–1427

Sprouffske, K., and Wagner, A. 2016. Growthcurver: An R package for obtaining interpretable metrics from microbial growth curves. BMC Bioinformatics. 17:17–20

Wetmore, K. M., Price, M. N., Waters, R. J., Lamson, J. S., He, J., Hoover, C. A., Blow, M. J., Bristow, J., Butland, G., and Arkin, A. P. 2015. Rapid quantification of mutant fitness in diverse bacteria by sequencing randomly bar-coded transposons. MBio. 6:1–15
